# The PMT-driven *p*-coumaroylation of poplar lignins impacts lignin structure and improves wood saccharification

**DOI:** 10.1101/2021.02.16.431462

**Authors:** Catherine Lapierre, Richard Sibout, Françoise Laurans, Marie-Claude Lesage-Descauses, Annabelle Déjardin, Gilles Pilate

## Abstract

Transgenic poplars (*Populus tremula x Populus alba*, clone INRA 717-1B4) were produced by introducing the *Brachypodium distachyon Bradi2g36910* (*BdPMT1*) gene driven by the Arabidopsis (*Arabidopsis thaliana)* Cinnamate 4-Hydroxylase (*AtC4H*) promoter in the wild-type (WT) line and in a line overexpressing the Arabidopsis Ferulate 5-Hydroxylase (*AtF5H). BdPMT1* encodes a transferase which catalyzes the acylation of monolignols by *p-*coumaric acid (CA). Several *BdPMT1*- OE/WT and *BdPMT1-OE/AtF5H-OE* transgenic lines were grown in the greenhouse and *BdPMT1* expression in xylem was confirmed by RT-PCR. The analysis of the cell walls (CW) of poplar stems and of corresponding purified dioxan lignins (DL) revealed that the *BdPMT1*-OE lignins were as *p*-coumaroylated as the lignins of C3 grass straws. For some transformants, CA levels even reached about 11 mg/g CW and 66 mg/g DL, which by far exceeds those of *Brachypodium* or wheat samples. This unprecedentedly high *p*-coumaroylation of poplar lignins affected neither the poplar growth, nor the stem lignin content. By contrast, the transgenic lignins were structurally modified, with an increase of terminal units with free phenolic groups. Relative to controls, this increase argues for a reduced polymerization degree of *BdPMT1*-OE lignins and makes them more soluble in cold NaOH solution. The *p*-coumaroylation of poplar samples, up to the levels of C3 grasses, improved the saccharification yield of alkali-pretreated poplar CW. These results establish that the genetically-driven *p*-coumaroylation of lignins is a promising strategy to make wood lignins more susceptible to the alkaline treatments that can be used during the industrial processing of lignocellulosics.

**One-sentence summary:** The expression of a grass p-coumaroyl-CoA:monolignol transferase induces a high p-coumaroylation of poplar lignins and a better saccharification of alkali-pretreated poplar wood without growth penalty

## INTRODUCTION

Wood appears as a major feedstock for traditional or innovative biorefineries producing pulp, chemicals or fermentable sugars. However, most industrial fractionations of lignocellulosics are detrimentally affected by lignins. For instance, the enzymatic hydrolysis of cellulose into glucose, referred to as saccharification, is severely hampered by lignins that hinder the accessibility of enzymes to CW polysaccharides. Indeed, the economically effective production of cellulosic ethanol necessitates costly, polluting and energy-intensive pretreatments that most often aim at reducing the lignin shield effect (Yang and Wyman, 2008; Sun et al., 2016). Since the last decades, lignin engineering in trees has been the subject of intensive studies to produce tailor-made wood more amenable to efficient deconstruction by milder processes (Pilate et al., 2012; Chanoca et al., 2019; Mahon and Mansfield, 2019). However, lignins play key roles in wood and sufficient lignin amounts are required to warrant tree growth, development and defense. On this basis, reducing lignin content may result in impaired tree growth and redesigning lignin structure appears as a better strategy to obtain wood biomass more adapted to industrial deconstruction without yield penalty.

Lignins primarily result from the enzymatically-driven oxidation of monolignols, mainly coniferyl alcohol and sinapyl alcohol that give rise to guaiacyl (G) and syringyl (S) units, respectively. It is now well established that lignin biosynthesis is very plastic and that, besides the main monolignols, a number of other molecules may participate to the formation of lignin polymers (Mottiar et al., 2016; del Río et al., 2020). For instance, *p*-coumaroylated sinapyl alcohol and, to a lower extent, *p*-coumaroylated coniferyl alcohol, are naturally incorporated into grass lignins (Grabber et al., 1996; Lu and Ralph, 1999; Hatfield et al., 2009; Ralph, 2010). This *p*-coumaroylation of grass monolignols is specifically catalyzed by a *p*-coumaroyl-coenzyme A monolignol transferase (PMT) studied in various grass species (Hatfield et al., 2009; Withers et al., 2012; Marita et al., 2014; Petrik et al., 2014). The *p*-coumaroylation of dicot lignins was recently achieved by introducing the rice *PMT* gene into poplar and arabidopsis plants (Smith et al., 2015), but the *p*-coumaroylation level of transgenic dicot CW reported in this study was modest (varying from 1 to 3.5 mg/g CW) and much lower than that of lignified grass stems (CA ranging from 6 to 39 mg/g CW) (Hatfield et al., 2009). By contrast, the introduction of two different *Brachypodium PMT* genes (*BdPMT1* or *BdPMT2*) under the control of the *AtC4H* promoter into various Arabidopsis lines boosted the *p*-coumaroylation of mature stem lignins up to the grass lignin level (Sibout et al., 2016). In addition to a high CA content, the Arabidopsis *BdPMT1-OE* lignins displayed other traits specific to grass lignins, *i.e.* a high frequency of free phenolic units in lignins and an increased solubility in cold alkali.

In this work, we explored the potential of introducing the *proAtC4H::BdPMT1* construct into poplar in order to beneficially tailor lignin structure without biomass penalty. To this end, *BdPMT1* was expressed not only in the poplar WT background, but also in a transgenic poplar line overexpressing the *AtF5H* gene (*AtF5H*-OE). By so doing, we obtained several independent transformants that were grown in the greenhouse together with the corresponding controls during 3 months. In this study, we first evaluated the growth of the *BdPMT1-OE* lines and the *p*-coumaroylation of their stem lignins, as compared to control trees. We then investigated the effect of the *BdPMT1* expression on lignin content and structure before subjecting the transgenic and control poplar stems to alkali-solubilization assays and saccharification tests.

## RESULTS AND DISCUSSION

### The Expression of Heterologous *BdPMT1* Gene under the Control of the *AtC4H* Promoter Does not Alter Poplar Growth

The *Bd*PMT1 acyltransferase (referred to as Bradi2g36910) has been shown to be specific to monolignol *p*-coumaroylation (Petrik et al., 2014). It is also the closest homologue of the rice *OsPMT* that was introduced by Smith et al. (2015) into poplar and Arabidopsis plants. As the *At*C4H promoter confers a vascular specific expression (Bell-Lelong et al., 1997), the *proAtC4H::BdPMT1* construct was introduced into poplar trees in order to preferentially express *BdPMT1* in the xylem tissues during the lignification step. The transformation was performed in two poplar genetic backgrounds, the WT line and a transgenic line overexpressing the *AtF5H* gene. The *AtF5H* expression was driven by a poplar cellulose synthase A4 promoter, known to be highly active in the fibers and vessels of poplar developing xylem (Hai et al., 2016). The *AtF5H-OE* poplar line was chosen to test the hypothesis that the *p*-coumaroylation of poplar lignins may be favored by a high frequency of S units based on the two following published data: a) the *p*-coumaroylation of grass lignins mostly occurs on S units (reviewed in Ralph, 2010; Karlen et al., 2018)) and b) overexpressing the *AtF5H* gene in poplar substantially increases the frequency of S lignin units (Franke et al., 2000).

The *Agrobacterium tumefaciens-mediated* transformation yielded 14 independent transformants in the WT background (referred to as *BdPMT1*-OE/WT lines) and 9 in the *AtF5H*-overexpressing background (referred to as *BdPMT1-OE/AtF5H-OE* lines). Three *BdPMT1-OE/WT* lines and five *BdPMT1-OE/AtF5H-OE* lines were selected for further analyses: they were acclimatized and grown for three months in the greenhouse together with corresponding control plants (Supplemental Fig.S1 A). RT-PCR with *BdPMT1* specific primers revealed a substantial *BdPMT1* transcript abundance in developing xylem, with some variations between the *BdPMT1-OE* lines, whereas no *BdPMT1* expression could be detected in the WT or *AtF5H-OE* control trees. Likewise, when using primers directed to *AtF5H*, a strong RT-PCR signal was observed in all the *AtF5H-OE* transgenic lines.

In the *BdPMT1* -OE/WT lines, the poplar plants did not show any phenotype different from the WT trees. However, in the *AtF5H-OE* background, two lines (referred to as lines 5 and 20.2) displayed patches of reddish coloration in the developing xylem, mostly at the nodes (Supplemental Fig. S1 B-C). Relative to the control trees, the *BdPMT1-OE* did not induce any significant difference in growth and height (Fig. 1).

**Figure 1.**
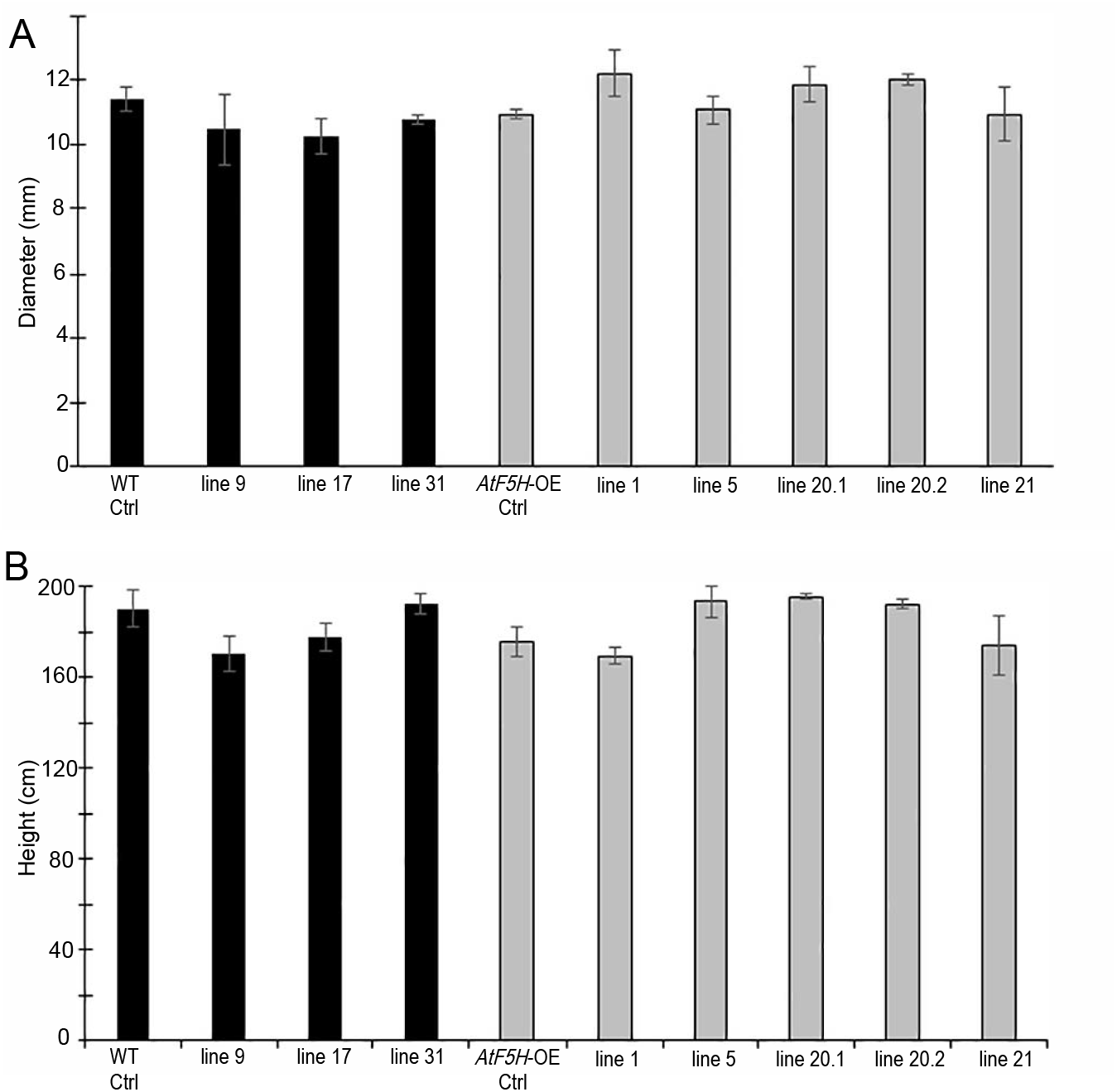
Growth response to the introduction of the *proAtC4H::BdPMT1* construct into the poplar WT background (black bars) and into the *AtF5H*-OE background (grey bars), as compared to control (Ctrl) trees. The basal diameter (A) and the tree height (B) were measured on 3-month-old greenhouse-grown trees. Data are means (and SD) of biological triplicates, except for lines 20.1 and 20.2 (biological duplicates).

### The *BdPMT1-OE* Poplar Stems and their Corresponding Purified Dioxane Lignin Fractions Are *p*-Coumaroylated up to the Levels of C3 Grass Samples

CW samples from the stems of 3-month-old greenhouse-grown poplar trees were subjected to mild alkaline hydrolysis to quantify *p*-hydroxybenzoic acid (Bz), *p*-coumaric acid (CA) and ferulic acid (FA) ester-linked to CW polymers. Poplar wood is typified by the occurrence of *p*-hydroxybenzoic acid ester-linked to lignins (Smith, 1955; Venverloo, 1969) and preferentially to the γ position of S lignin units (Lu et al., 2004; Morreel et al., 2004). Most *BdPMT1-OE* poplar samples displayed similar *p*-hydroxybenzoylation levels as their corresponding controls (Table I). In addition to Bz, mild alkaline hydrolysis of poplar samples released small amounts of FA consistently obtained in slightly smaller quantities in *BdPMT1*-OE/WT lines compared to WT, whereas *BdPMT1-OE/AtF5H-OE* lines 5 and 20.2 delivered more FA than the other *AtF5H-OE* lines (Table I). In plant CW, FA preferentially acylates non cellulosic polysaccharides (Ishii, 1997) and the small differences of FA esters between poplar lines might reflect some structural variations in these CW components.

**Table I.** Amount of *p*-hydroxybenzoic acid (Bz), *p*-coumaric acid (CA) and ferulic acid (FA) released by mild alkaline hydrolysis of extract-free poplar stems from *BdPMT1-OE* lines obtained in the WT and *AtF5H-OE* backgrounds, as compared to their respective controls. The data represent mean values (and SD) from n biological replicates. asterisks denote significant differences (one-way ANOVA) compared to the value of the corresponding control (*: P < 0.05; **: P < 0.001).

| Line (n replicates) | Bz<br>mg/g CW | CA<br>mg/g CW | FA<br>mg/g CW |
| --- | --- | --- | --- |
| WT control (3) | 3.86 (0.10) | 0.01 (0.00) | 0.22 (0.00) |
| *BdPMT1*-OE/WT line 9 (3) | 3.21 (0.51) | 7.12 (0.49)** | 0.15 (0.02)* |
| *BdPMT1*-OE/WT line 17 (3) | 3.66 (0.28) | 10.69 (0.49)** | 0.18 (0.01)* |
| *BdPMT1*-OE/WT line 31 (3) | 3.57 (0.11) | 3.63 (0.42)** | 0.10 (0.00)* |
| *AtF5H*-OE control (4) | 3.29 (0.14) | 0.01 (0.00) | 0.06 (0.00) |
| *BdPMT1*-OE/*AtF5H*-OE line 1 (4) | 3.06 (0.56) | 0.76 (0.04)* | 0.07 (0.01) |
| *BdPMT1*-OE/*AtF5H*-OE line 5 (4) | 3.47 (0.26) | 11.06 (0.63)** | 0.31 (0.04)* |
| *BdPMT1*-OE/*AtF5H*-OE line 20.1 (2) | 3.66 (0.12) | 4.41 (0.18)** | 0.13 (0.02) |
| *BdPMT1*-OE/*AtF5H*-OE line 20.2 (2) | 3.53 (0.03) | 11.11 (0.10)** | 0.40 (0.07)* |
| *BdPMT1*-OE/*AtF5H*-OE line 21 (3) | 4.35 (0.08)* | 4.92 (0.19)** | 0.13 (0.02) |

While expressing the *BdPMT1* gene both in the WT and the *AtF5H-OE* backgrounds had little or no effect on Bz or FA units ester-linked to poplar CW, this transformation increased CW *p*-coumaroylation to different levels with CA quantity ranging between 0.76 to 11.11 mg/g CW, as compared to the trace amounts of the controls (Table I). Remarkably enough, this quantity was boosted up to about 11 mg/g CW in *BdPMT1*-OE/WT line 17, *BdPMT1-OE/AtF5H-OE* line 5 and *BdPMT1*- OE/*’AtF5H-OE* line 20.2. As compared to grass mature stems, the CA levels of these three poplar lines exceeded those of most C3 grass CW, but remained lower than those of C4 grass CW (Supplemental Table S1). With the exception of one line, the obtained *BdPMT1* -OE poplar lines were as *p*-coumaroylated as extract-free *proAtC4H::BdPMT1* Arabidopsis mature stems (CA amounts ranging between 3.5 and 12.6 mg/g CW) (Sibout et al., 2016). By contrast, these levels were much higher than the values reported for *OsPMT-OE* poplar lines (CA range: 1.2-3.5 mg/g CW) or for *OsPMT-OE* Arabidopsis lines (CA range: 1.0-2.0 mg/g CW) when *OsPMT* expression was driven by the CAULIFLOWER MOSAIC VIRUS promoter or by the CELLULOSE SYNTHASE7 promoter (Smith et al., 2015). In agreement with this study (Smith et al., 2015), the *p*-coumaroylation of poplar CW did not affect their *p*-hydroxybenzoylation (Table I). The high *p*-coumaroylation of poplar CWs obtained in the present work is very likely related to the efficiency of the *AtC4H* promoter, in agreement with recent data obtained with *BdPMT1* -OE Arabidopsis lines (Sibout et al., 2016).

Isolation of DL fractions followed by their mild alkaline hydrolysis recently proved to be an efficient strategy to demonstrate that CA units introduced in *BdPMT1* -transformed arabidopsis plants are ester-linked to lignins (Sibout et al., 2016). The isolation method consists in mild acidolysis (refluxing CW samples in dioxane/0.2 M aq. HCl for 30 min under N_2_), which provides a rough lignin extract then purified to recover DL fractions. This isolation method relies on the hydrolysis of some ether bonds in lignins to make the insoluble native lignin polymers partially soluble into the reaction medium. The purified DL fractions contain a low amount of sugar contaminants (< 10% by weight) and the mild isolation procedure mostly preserve lignin-linked CA esters, if present (Chazal et al., 2014). Purified poplar DL fractions were isolated from a few control and *BdPMT1-OE* poplar lines and then subjected to mid-IR spectroscopy. Their mid-IR spectra not only confirmed their low contamination by sugar components, but also suggested that the lignin fractions isolated from *BdPMT1* -OE*/*WT and *BdPMT1-OE/AtF5H-OE* lines were enriched in CA esters (Supplemental Fig. S2). Relative to their respective controls, the IR spectra from *BdPMT1-OE* lines displayed increased signals at 1604, 1164 and 833 cm^-1^, which can be assigned to the occurrence of CA units (Chazal et al., 2014). More importantly, high CA amounts (from 31 to 66 mg/g DL, Table II) were released by mild alkaline hydrolysis of the purified DL fractions isolated from *BdPMT1-OE* poplar lines, as confirmed by both HPLC and GC/MS analyses (Supplemental Fig. S3). The upper values were similar to the CA levels of DL fractions isolated from C3 grass CW, but remained lower than those of DL fractions isolated from C4 grass species (Supplemental Table S1). Alkaline hydrolysis of the DL fractions isolated from control samples released noticeable amounts of CA units (Table II), which reveals that CA acylates poplar lignins to a weak extent and in agreement with results obtained for Arabidopsis lignins (Sibout et al., 2016). The CA contents of DL fractions from *BdPMT1-OE* poplar line were found to be 6-to 10-fold higher than those from the corresponding CW. Such an outstanding enrichment definitely establishes that most CA units introduced in the transgenic poplars are ester-linked to lignins.

**Table II.** Amount of *p*-coumaric acid (CA) released by mild alkaline hydrolysis of DL fractions isolated from control and *BdPMT1-OE* lines obtained in the WT and *AtF5H-OE* backgrounds. The data represent mean values (and SD) from technical duplicates

| Line | CA<br>mg/g DL |
| --- | --- |
| WT control | 3.21 (0.17) |
| *BdPMT1*-OE/WT line 9 | 50.00 (0.77) |
| *BdPMT1*-OE/WT line 17 | 66.52 (0.47) |
| *BdPMT1*-OE/WT line 31 | 31.36 (0.43) |
| *AtF5H*-OE control | 0.87 (0.04) |
| *BdPMT1*-OE/*AtF5H*-OE line 5 | 66.43 (1.43) |
| *BdPMT1*-OE/*AtF5H*-OE line 21 | 33.98 (0.53) |

### Analytical Pyrolysis Further Confirms the High *p*-Coumaroylation of *BdPMT1*- OE Poplar Lines

The main advantages of the pyrolysis-gas chromatography/mass spectrometry (Py-GC/MS) method is its high throughput screening capabilities together with its low sample demand (Ralph and Hatfield, 1991; Lapierre, 1993). When subjected to this method, lignified CW samples provide lignin-derived phenolics originating from G and S lignin units. In addition, during pyrolysis, ester-linked Bz and CA units (if present) are decarboxylated to produce phenol (P) and vinylphenol (VP), respectively. The relative abundances (area %) of the main G and S pyrolysis products and of P and VP generated from the poplar CW samples are listed in Table III. The relative percentage of pyrolysis-derived P did not discriminate the various transgenic samples from their control. This result is quite consistent with mild alkaline hydrolysis which provided similar Bz amounts from most transgenic lines and their respective controls. By contrast, the relative importance of VP was dramatically increased in the *BdPMT1*-OE lines as compared to their controls. Such a relative increase concomitantly decreased the relative percentage of the lignin-derived pyrolysis compounds ((S+G) in Table III). Even though the pyrolysis VP might originate from tyrosine residues of putatively present protein contaminants, it is essentially produced from the decarboxylation of CW-linked CA units (Ralph and Hatfield, 1991). The VP relative abundance was found to nicely echoe the level of alkali-releasable CA, as revealed by the positive correlation between CA amount and the % VP (R^2^ = 0.982, Supplemental Fig. S4). In other words, the relative importance of pyrolysis-derived VP may be viewed as a good signature of the CW *p*-coumaroylation level. To further confirm that VP prominently originates from CA decarboxylation, a few pyrolysis assays were carried out in the presence of tetramethylammonium hydroxyde (TMAH). The TMAH-Py-GC/MS method yields methyl 4-methoxybenzoate (Bz_Me_) and methyl 4-methoxy-*p*-coumarate (CA_Me_) from Bz and CA units, respectively (Kuroda et al., 2001; Kuroda et al., 2002). As shown in the pyrograms oulined in Fig. 2, the relative intensity of the Bz_Me_ peak was similar in the *BdPMT1-OE* and in their corresponding controls whereas the CAMe peak was prominent in the *BdPMT*-OE poplar lines.

**Figure 2.**
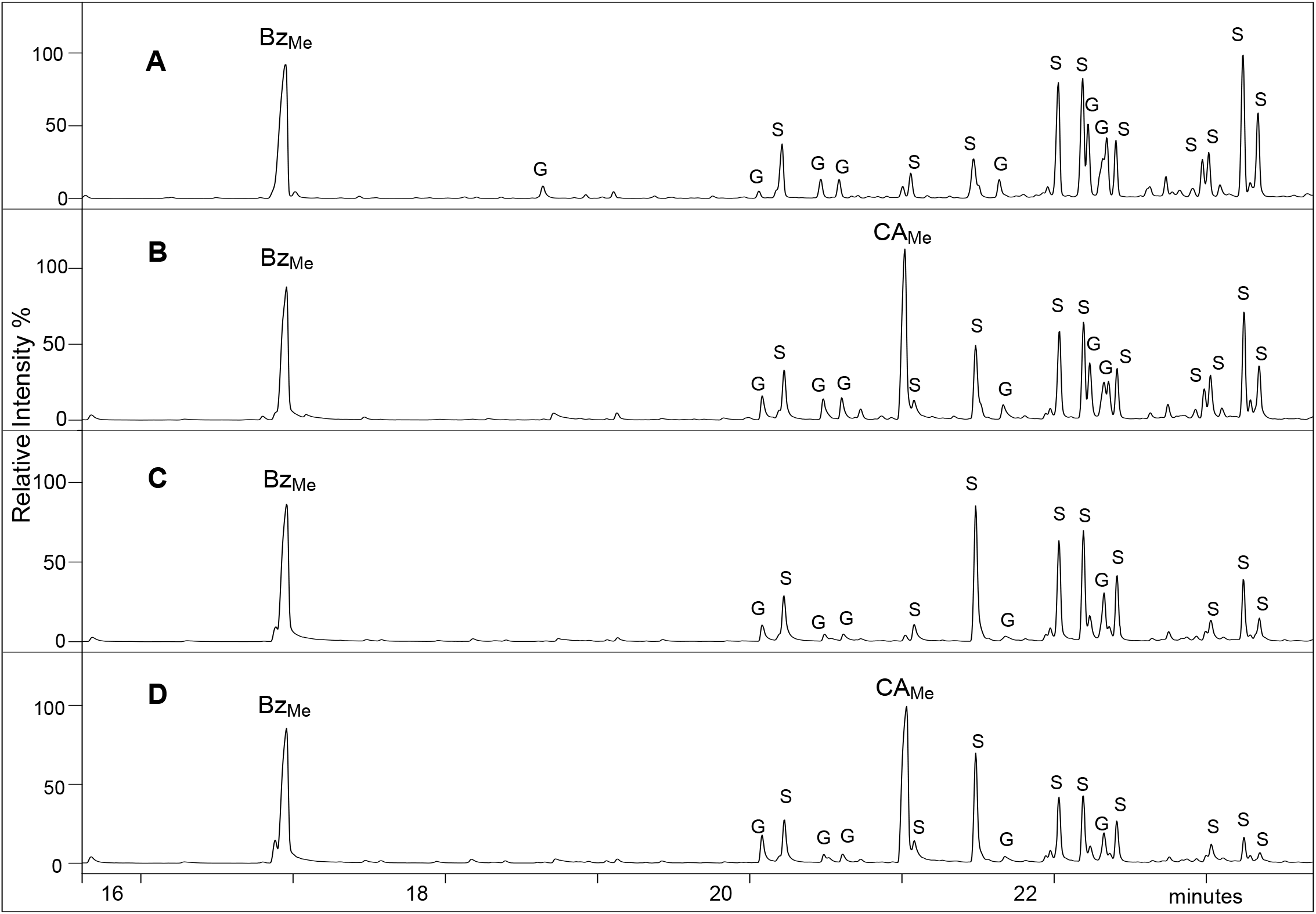
TMAH-Py-GC/MS traces of poplar CW from A) WT control, B) *BdPMT1*-OE/WT line 9, C) *AtF5H*-OE control and D) *BdPMT1-OE/AtF5H-OE* line 5. Bz_Me_: 4-methoxybenzoate; CA_Me_: methyl 4-methoxy-*p*-coumarate; peaks quoted G and S correspond to methylated G and S compounds, respectively.

**Table III.** Relative percentage values of the peaks assigned to the main phenolics released by Py-GC/MS of poplar CW from *BdPMT1-OE* lines obtained in the WT and *AtF5H-OE* backgrounds, as compared to their respective controls. These area values are expressed as percentage of the total area per sample (set to 100). The data represent mean values (and SD) from n biological replicates. asterisks denote significant differences (one-way ANOVA) compared to the value of the corresponding control (*: P < 0.05; **: P < 0.01).

| Line (n replicates) | Phenol | 4-vinylphenol | (S+G) compounds<sup>a</sup> | S/G ratio |
| --- | --- | --- | --- | --- |
| WT control (4) | 5.33 (0.46) | 0.21 (0.07) | 94.47 (0.41) | 2.82 (0.21) |
| *BdPMT1*-OE/WT line 9 (4) | 4.60 (0.94) | 10.73 (0.35)** | 84.67 (1.22)** | 2.88 (0.13) |
| *BdPMT1*-OE/WT line 17 (4) | 5.43 (0.43) | 15.44 (0.71)** | 79.13 (0.57)** | 2.83 (0.12) |
| *BdPMT1*-OE/WT line 31 (4) | 5.06 (0.56) | 5.40 (1.46)** | 89.54 (1.72)** | 2.72 (0.19) |
| *AtF5H*-OE control (4) | 4.99 (0.70) | 0.10 (0.04) | 95.01 (0.70) | 4.11 (0.29) |
| *BdPMT1*-OE/*AtF5H*-OE line 1 (4) | 4.40 (1.17) | 1.15 (0.010)* | 94.47 (1.23) | 4.32 (0.52) |
| *BdPMT1*-OE/*AtF5H*-OE line 5 (4) | 4.92 (0.68) | 15.01 (0.86)** | 80.08 (0.67)** | 2.73 (0.24)* |
| *BdPMT1*-OE/*AtF5H*-OE line 20.1 (2) | 4.99 (0.25) | 6.65 (0.30)** | 88.37 (0.55)** | 4.27 (0.12) |
| *BdPMT1*-OE/*AtF5H*-OE line 20.2 (2) | 4.51 (1.03) | 16.42 (1.52)** | 79.08 (0.48)** | 2.50 (0.16)* |
| *BdPMT1*-OE/*AtF5H*-OE line 21 (3) | 6.21 (0.42) | 7.43 (0.25)** | 86.35 (0.52)** | 4.27 (0.24) |

<sup>a</sup> G compounds include: guaiacol, 4-methylguaiacol, 4-ethylguaiacol, 4-vinylguaiacol, 4-allylguaiacol (2 isomers), vanillin, acetoguaiacone, guaiacylacetone; S compounds include: syringol, 4-methylsyringol, 4-ethylsyringol, 4-vinylsyringol, 4-allylsyringol (2 isomers), syringaldehyde, acetosyringone, syringylacetone.

The pyrolysis S/G ratio calculated from the relative importance of lignin-derived S and G pyrolysis compounds was not significantly affected in the *BdPMT1*-OE/WT lines (Table III). This result suggests that the proportion of G and S lignin unit is not affected by the transformation. In agreement with literature data (Franke et al., 2000; Stewart et al., 2009), this ratio was substantially increased in the *AtF5H-OE* control line as well as in the *BdPMT1-OE/AtF5H-OE* lines 1,20.1 and 21. By contrast, the *BdPMT1-OE/AtF5H-OE* lines 5 and 20.2, which were provided with a patchy reddish xylem coloration and the highest CA levels, displayed much lower pyrolysis S/G ratios (Table III), in agreement with the reduced Maüle staining observed on stem transverse sections from these lines (Supplemental Fig. S1 D). This result suggests that the substantial participation of *p*-coumaroylated monolignols to lignification somehow counteracted the *AtF5H-OE* related enrichment in S units.

### The *proAtC4H::BdPMT1* Transformation Has no or Little Effect on the Lignin Content of Poplar Stems, but a Strong Impact on Lignin Structure

The most *p*-coumaroylated transgenic poplar lines were analyzed for their lignin contents, using both the Klason Lignin (KL) and the Acetyl Bromide Lignin (ABL) methods (Table IV). With the exception of the *BdPMT1-OE/AtF5H-OE* line 5 displaying slightly higher KL and ABL contents, the *BdPMT1* transformation had no impact on the lignin content of the poplar stem CW. This result contrasts with those obtained for *proAtC4H::BdPMT1* Arabidopsis transformants provided with similar *p*- coumaroylation levels as these poplar transgenics, but with 10 to 30% lower lignin contents than their controls (Sibout et al., 2016). Introducing the *proAtC4H::BdPMT1* into Arabidopsis plants seemed to affect the metabolic flux to lignins and thereby the stem lignin content whereas such an effect was not observed in the *BdPMT1-OE* poplar lines.

**Table IV.**
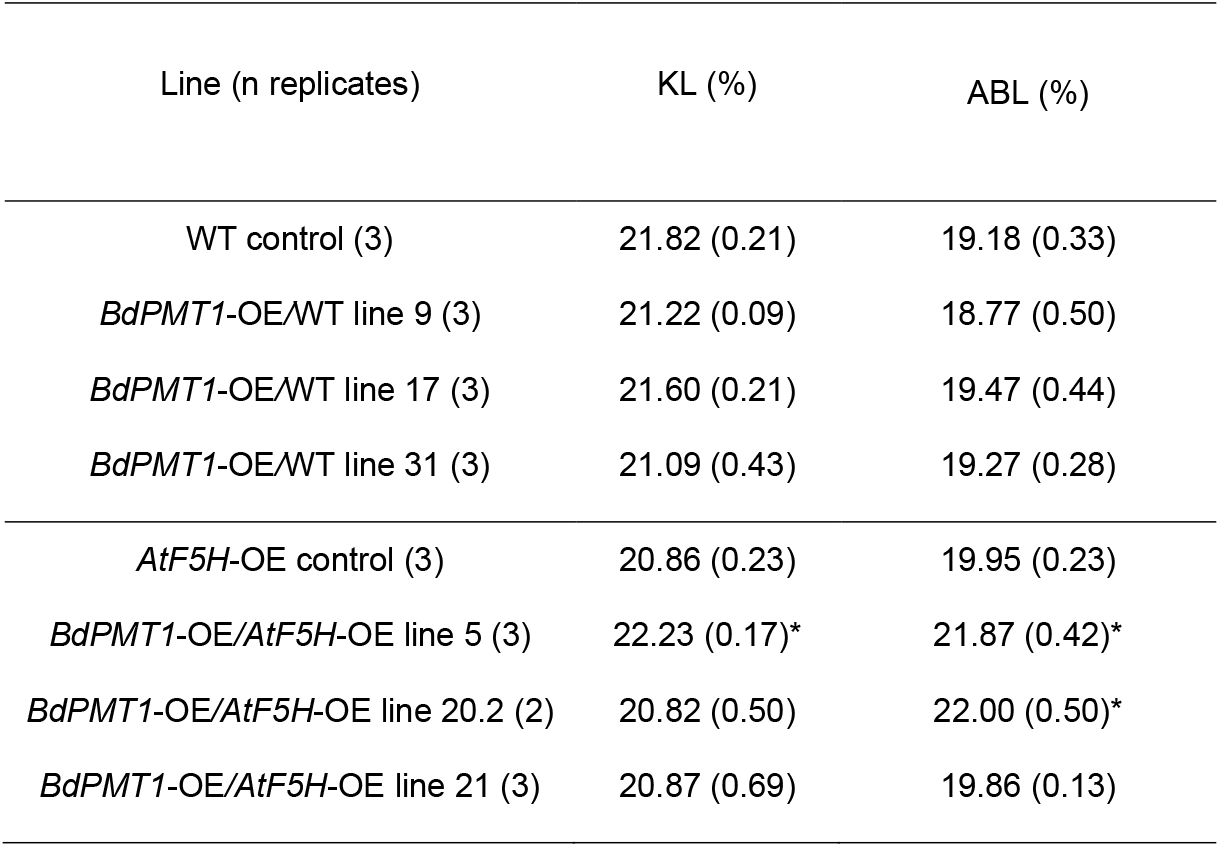
Lignin content of extract-free poplar stems from *BdPMT1-OE* lines obtained in the WT and *AtF5H*-OE backgrounds, as compared to their respective controls. The lignin content is expressed as weight percentage of the sample and was determined using the Klason Lignin (KL) and the Acetyl Bromide Lignin (ABL) methods The data represent mean values (and SD) from n biological replicates. asterisks denote significant differences (one-way ANOVA) with the control at P < 0.05.

A major structural trait of native lignins is their percentage of free phenolic groups, which has a strong impact on lignin susceptibility towards industrial alkaline or oxidative treatments. When thioacidolysis is performed on CW exhaustively permethylated with diazomethane or trimethylsilyldiazomethane (TMSD), the percentages of free phenolic groups in β-O-4 linked G or S lignin units, referred to as % GOH or % SOH, can be evaluated. These percentages have been shown to nicely parallel that of the whole polymer (Lapierre, 2010). With the objective to evaluate the impact of the *BdPMT1* transformation on the structure of poplar native lignins, we employed this analytical approach, the principle of which is outlined in Fig. 3. Past studies have shown that the thioacidolysis yield is not affected by the mild permethylation procedure (Lapierre et al., 1988; Lapierre, 2010). Whatever the sample, the *p*-hydroxyphenyl (H) thioacidolysis monomers were found to be obtained as trace components (less than 1% of the monomer yield) and, in consequence, these minor H units were not considered in the following. In agreement with the Py-GC/MS data, the thioacidolysis S/G ratio was not affected by the *BdPMT1* transformation in the WT background (Table V). Not unexpectedly and as compared to the WT, the thioacidolysis S/G ratio was found to be drastically increased in the *AtF5H-OE* control sample as well as in the *BdPMT1-OE/AtF5H-OE* lines 20.1 and 21 (Table V). Consistently with the pyrolysis data (Table III), the two *BdPMT1-OE/AtF5H-OE* lines 5 and 20.2, which are provided with the highest *p*-coumaroylation levels (Table I), displayed S/G ratios close to those observed in the WT background. This result confirms that the high *p*-coumaroylation level of these two poplar lines somehow hinders the *AtF5H*-driven enrichment in S units by a mechanism which remains to be explained.

**Figure 3.**
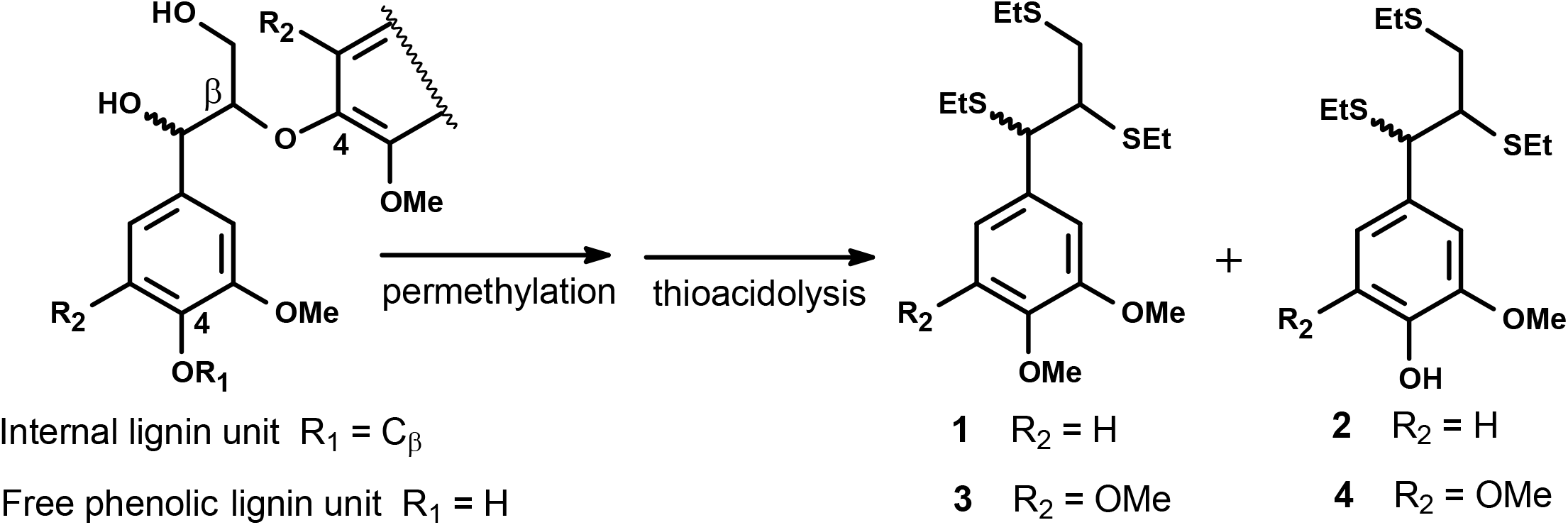
Principle of the evaluation of free phenolic units in lignin by thioacidolysis of permethylated samples. Lignin units only involved in β-O-4 bonds give rise to thioacidolysis guaiacyl (R_2_ = H) and syringyl (R_2_ = OMe) monomers. Terminal G and S units with free phenolic group (R_1_ = H) are first methylated at C4, then degraded to monomers **1** and **3** (erythro/threo mixture), respectively. Internal G and S units (R_1_ = C_β_ of another lignin sidechain) are degraded to monomers **2** and **4**, respectively (erythro/threo mixture).

**Table V.** Thioacidolysis of TMSD-methylated poplar CW from from *BdPMT1-OE* lines obtained in the WT and *AtF5H-OE* backgrounds, as compared to their respective controls. The S/G molar ratio corresponds to the ratio of the S monomers (**3** + **4**) to the G monomers (**1** + **2**) (monomers shown in Figure 3). The molar % of free phenolic groups in β-O-4 linked G or S units, referred to as % GOH or % SOH, is calculated according to the outlined formula. The data represent mean values (and SD) from n biological replicates. asterisks denote significant differences (one-way ANOVA) compared to the value of the corresponding control (*: P < 0.05; **: P < 0.01).

| Line (n replicates) | S/G molar ratio<br>(**3 + 4**)/(**1 + 2**) | % free phenolic units in βO-4 linked | |
| --- | --- | --- | --- |
|  |  | G or S units |  |
|  |  | % GOH | % SOH |
| | | $100 \times 1/(1+2)$ | $100 \times 3/(3+4)$ |
| WT control (3) | 2.05 (0.03) | 19.45 (0.22) | 2.81 (0.07) |
| *BdPMT1*-OE/WT line 9 (3) | 2.09 (0.18) | 22.85 (0.12)** | 3.65 (0.19)** |
| *BdPMT1*-OE/WT line 17 (3) | 2.12 (0.19) | 23.65 (0.48)** | 4.44 (0.25)** |
| *BdPMT1*-OE/WT line 31 (3) | 2.08 (0.06) | 21.09 (0.26)** | 3.44 (0.10)* |
| *AtF5H*-OE control (3) | 3.12 (0.13) | 20.85 (0.44) | 3.26 (0.03) |
| *BdPMT1*-OE/*AtF5H*-OE line 5 (3) | 1.84 (0.15)** | 22.66 (0.02)** | 4.34 (0.08)** |
| *BdPMT1*-OE/*AtF5H*-OE line 20.1 (2) | 3.36 (0.04) | 22.55 (0.59)* | 3.90 (0.09)* |
| *BdPMT1*-OE/*AtF5H*-OE line 20.2 (2) | 1.83 (0.02)** | 22.79 (0.22)** | 4.15 (0.19)** |
| *BdPMT1*-OE/*AtF5H*-OE line 21 (3) | 2.96 (0.15) | 23.10 (0.49)* | 4.02 (0.21)* |

In agreement with literature data (Lapierre, 1993, 2010), the control poplar samples displayed a % GOH and a % SOH close to 20% and 3%, respectively, which confirms that S units essentially are internal units. These percentages were significantly increased in the *p*-coumaroylated lignins of the *BdPMT1-OE* poplar lines (Table V). The increase in % GOH or in % SOH was found to be nicely correlated to the CA level of the *BdPMT1*-OE/WT lines (R^2^ = 0.95 for % GOH and 0.93 for % SOH) (Fig. 4). This result means that the incorporation of *p*-coumaroylated monolignols in poplar lignins increases the frequency of free phenolic terminal units relative to internal units. Such a structural change may be accounted for by the occurrence of lignin polymers with lower polymerization degree and/or with a higher content of biphenyl or biphenyl ether branching structures.

**Figure 4.**
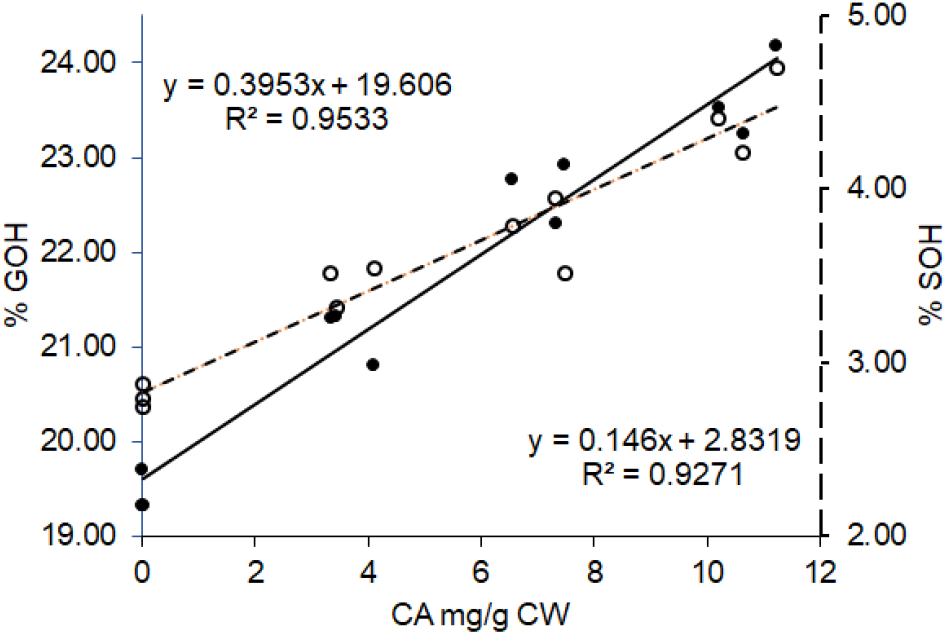
Relationships between the CA amounts in poplar CW and the percentage of G lignin units with free phenolic groups (% GOH, black circles, full line) or the percentage of S lignin units with free phenolic groups (% SOH, white circles, dotted lines). The lignin structural traits % GOH and % SOH are evaluated by thioacidolysis of permethylated samples for *BdPMT1-OE* poplars and their WT controls.

The alkaline hydrolysis of the DL fractions isolated from the *BdPMT1-OE* poplar lines revealed that their CA units were primarily ester-linked to lignins. With the objective to more precisely localize these CA esters on lignin units, we subjected some poplar samples to 1-hour long thioacidolysis experiments, followed by Raney nickel desulfuration in order to identify the syringylpropanol and/or guaiacylpropanol units acylated by *p*-dihydrocoumaric acid (diHCA). This short thioacidolysis time is necessary as CA esters do not survive the standard 4-hour long thioacidolysis method (Lapierre, 1993; Sibout et al., 2016). When applied to the *BdPMT1* -OE*/*WT line 17, the method provided substantial amount of syringylpropanol acylated by diHCA while this dimer could not be observed with a longer thioacidolysis duration (Supplemental Fig. S5 and Supplemental Table S2). Interestingly enough and by contrast to the results reported by Smith et al. (2015), its G analogue could not be detected. Taken together and similarly to grass lignins, these results support the hypothesis that the *p*-coumaroylation of poplar transformants primarily involves S lignin units.

The analysis of the lignin-derived dimers obtained with the standard thioacidolysis method followed by Raney nickel desulfuration confirmed that lignins from *BdPMT1-OE* poplars were structurally different from control lignins. The main difference was related to the relative importance of the syringaresinol-derived dimer, expressed as percentage of the total area of the main dimers (set to 100): this relative percentage was increased from (18.5 ± 1.7)% for the WT sample up to (26.1 ± 1.9)% for the *BdPMT1* -OE/WT line 17 line (mean and SD for duplicate analyses). The syringaresinol structures exclusively originate from the dimerization of sinapyl alcohol and are thus starting points for lignin growth (Ralph et al., 2004). Their higher relative recovery from the *BdPMT1*-OE/WT line 17 further argues for the occurrence of lignin polymers with lower polymerization degrees than in the control sample.

### The *BdPMT1*-driven Substantial *p*-Coumaroylation of Poplar Samples Makes their Lignins more Easily Solubilized in Cold Alkali

The enrichment in free phenolic G and S units is very likely to improve the lignin susceptibility to alkaline treatments that are employed in chemical pulping or in the cellulose-to-ethanol conversion process. The impact of % GOH on the CW delignification induced by alkaline treatment has been established for a long time for grass samples (Lapierre et al., 1989; Lapierre, 2010) and confirmed for poplar trees deficient in Cinnamyl Alcohol Dehydrogenase (CAD) activity (Lapierre et al., 1999; Lapierre et al., 2004; Van Acker et al., 2017), for tobacco plants deficient in Cinnamoyl-Coenzyme A Reductase (CCR) activity (O’Connell et al., 2002) and for *BdPMT1* -transformed Arabidopsis lines (Sibout et al., 2016). The results of a mild alkaline treatment applied to the poplar samples are shown in Table VI. The residue recovered after this treatment, referred to as the saponified residue (SR), was obtained with similar yields whatever the line. However, its lignin amount was found to be lower in *BdPMT1-OE* lines relative to their control (Table VI). Consistently with these results, the percentage of the alkali-soluble lignin (%Alk-L) revealed that the *BdPMT1-OE* lines are more easily delignified by the employed mild alkaline treatment. Whereas 15 to 20 % of the lignin polymers were solubilized by cold alkali for the controls, the %Alk-L was substantially increased in the *BdPMT1-OE* lines. As reported for transgenic CAD-or CCR-deficient plants (Lapierre et al., 1999; O’Connell et al., 2002), increasing the % GOH has beneficial effects on the kraft pulping properties of the lignocellulosic biomass, thereby decreasing the energy and environmental costs of this industrial process.The introduction of *BdPMT1* in trees would likely improve the pulping properties of poplar wood.

The relationship of the free phenolic groups in poplar lignins to their susceptibility towards cold alkaline treatment is further illustrated in Fig. 5. On this scheme, we have gathered the data from 17 different poplar lines, comprising the current *BdPMT1-OE/WT* lines and CAD-deficient ones (Lapierre et al., 2004), together with their respective controls. The effect of the % GOH structural property onto the solubility of poplar lignins in cold alkali is supported by the positive correlation between % GOH and % Alk-L (R^2^ = 0.9513).

**Figure 5.**
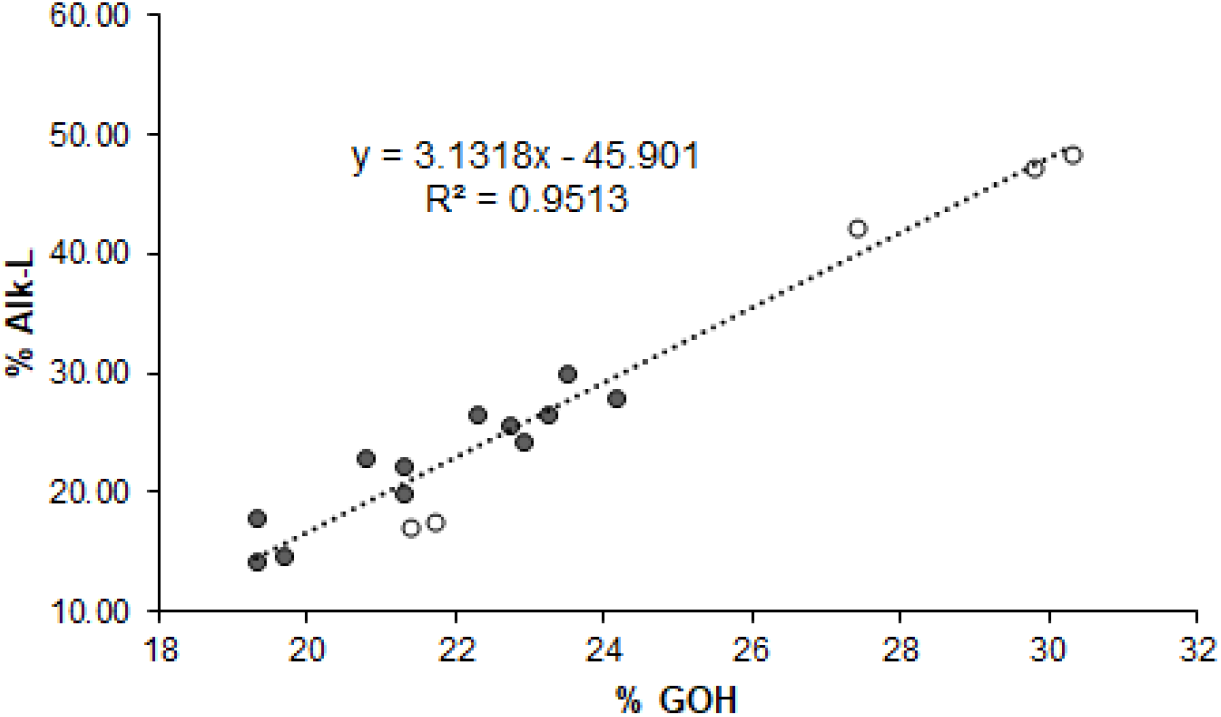
Relationship between the percentage of G lignin units with free phenolic groups (% GOH) and the solubility of poplar lignins in cold alkali (% Alk-L). The data correspond to *BdPMT1-OE* trees and their WT controls (black circles) as well as to CAD-deficient trees and their corresponding controls (white circles).

### The BdPMT1-Driven *p*-Coumaroylation of Poplar Samples Results in Improved Saccharification after Cold Alkaline Pretreatment

It is well established that the detrimental role of lignins on the cost-effective enzymatic conversion of lignocellulosic polysaccharides into fermentable sugars makes necessary the use of pretreatments (Yang and Wyman, 2008; Wang et al., 2015; Sun et al., 2016). From the analytical data that we obtained so far on the *BdPMT1-OE* poplar lines, we could anticipate that an alkaline pretreatment would be well suited to reduce the lignin-related recalcitrance of poplar wood to saccharification. Accordingly, the saccharification experiments run on the poplar samples were preceded by a cold alkaline pretreatment (aq. NaOH 1M, overnight, room temperature). The saccharification efficiency was evaluated both by the weight loss (% WL) and by the amount of released glucose (Glc) (Table VII). Both parameters were higher in the *BdPMT1-OE* lines, compared with their control. Not unexpectedly and within each background, the best saccharification results were obtained for the lines provided with the concomitantly highest CA level, % GOH and % Alk-L. The enrichment of poplar lignins in free phenolic groups made these lignins more easily solubilized in alkali, which consequently improved the saccharification of alkali-pretreated samples. Taken together, these results reveal that the lignins from the current *BdPMT1-OE* poplar plants share common features with grass lignins. As compared to non grass lignins from WT plants, these common features are a) a substantial *p*-coumaroylation of S lignin units, b) a higher level of free phenolic units, and c) a higher solubility in cold alkali. At this point, we may hypothesize that, similar to grass lignins, lignins from the *BdPMT1-OE* poplar lines obtained herein are distributed in the cell walls as small lignin domains which are both rich in free phenolic groups and more easily extracted by cold alkali treatment (Lapierre, 2010).

**Table VI.** Impact of a mild alkaline treatment (aq. NaOH 1 M, overnight, room temperature) on extract-free poplar stems from control and *BdPMT1-OE* lines obtained in the WT and *AtF5H*-OE backgrounds. The percentage of the recovered saponified residue (% SR) is expressed relative to the initial sample. The lignin content of the SR sample is measured as acetyl bromide lignin (% ABL). The percentage of alkali-soluble lignins (% Alk-L) is calculated from the ABL content of the CW and from the % SR recovery yield. The data represent mean values (and SD) from biological triplicates. asterisks denote significant differences (one-way ANOVA) compared to the value of the corresponding control (*: P < 0.05; **: P < 0.01).

| Line | % SR | % ABL in SR | % Alk-L |
| --- | --- | --- | --- |
| WT control | 68.03 (0.88) | 23.69 (0.04) | 15.5 (2.1) |
| *BdPMT1*-OE/WT line 9 | 67.17 (0.46) | 20.85 (0.47)** | 25.5 (1.1)** |
| *BdPMT1*-OE/WT line 17 | 65.25 (0.73)* | 21.52 (0.24)** | 28.1 (1.7)** |
| *BdPMT1*-OE/WT line 31 | 68.17 (0.22) | 22.11 (0.75)* | 21.7 (1.6)* |
| *AtF5H*-OE control | 69.02 (0.65) | 22.96 (0.28) | 20.5 (1.3) |
| *BdPMT1*-OE/*AtF5H*-OE line 5 | 66.89 (0.80) | 21.05 (0.43)** | 36.1 (1.1)** |
| *BdPMT1*-OE/*AtF5H*-OE line 21 | 67.65 (1.28) | 21.52 (0.13)* | 26.4 (1.7)* |

**Table VII.** Saccharification of the poplar saponified residues (SR) obtained after a mild alkaline treatment (aq. NaOH 1 M, overnight, room temperature) and corresponding to *BdPMT1-OE* lines obtained in the WT and *AtF5H-OE* backgrounds, as compared to their respective controls. The saccharification efficiency is evaluated both by the weight loss (%WL) and by the released glucose (Glc). The data represent mean values (and SD) from biological triplicates. asterisks denote significant differences (one-way ANOVA) compared to the value of the corresponding control (*: P < 0.05; **: P < 0.01).

| SR from Line | % WL | Glc<br>mg.g<sup>-1</sup> SR |
| --- | --- | --- |
| WT control | 39.8 (1.3) | 307.7 (16.0) |
| *BdPMT1*-OE/WT line 9 | 52.1 (1.6)** | 417.5 (23.2)** |
| *BdPMT1*-OE/WT line 17 | 55.1 (2.0)** | 452.4 (17.7)** |
| *BdPMT1*-OE/WT line 31 | 45.8 (0.3) | 369.3 (17.9)* |
| *AtF5H*-OE control | 44.2 (2.2) | 401.4 (6.7) |
| *BdPMT1*-OE/*AtF5H*-OE line 5 | 56.4 (1.3)** | 510.1 (9.1)** |
| *BdPMT1*-OE/*AtF5H*-OE line 21 | 49.4 (1.5)** | 461.7 (22.4)** |

## CONCLUSION

In this study, we have shown that *p*-coumaroylating poplar lignins up to the level of grass lignins has consequences that go far beyond a simple lignin decoration and that deeply change not only lignin structural traits, but also important industrial potentialities of lignified CW. Remarkably enough the *proAtC4H::BdPMT1* transformation introduced neither any growth penalty, nor reduced lignin content in the various transgenic greenhouse-grown poplar lines that were obtained in two genetic backgrounds. In agreement with a recent study (Sibout et al., 2016), choosing the lignin-specific *AtC4H* promoter to drive the heterologous expression of *BdPMT1* in dicot CW had very likely a key role in changing wood properties.

Since the last decades and with the objective to facilitate the industrial conversion of lignocellulosics into pulp or into bioethanol, many approaches have been used to genetically modify lignin content and/or structure (reviewed in (Boerjan and Ralph, 2019; Halpin, 2019; Mahon and Mansfield, 2019; Ralph et al., 2019)). Among the lignin structural traits that can be affected by the genetic transformation of angiosperm species, the S/G ratio is probably the most systematically scrutinized one (Chanoca et al., 2019). By contrast, the relative frequency of free phenolic units in native lignins is a key structural trait which is surprisingly overlooked despite its biological significance and its major effect on the susceptibility of lignins to alkaline or oxidative treatments. In past studies, redesigning native lignins with more free phenolic groups (and therefore with increased alkali-solubility) could be obtained with other genetic transformations, such as CCR or CAD down-regulation (O’Connell et al., 2002; Lapierre et al., 2004). In this work, we provide another compelling evidence that the genetically-driven increase of free phenolic units in lignins is an efficient strategy for the rational design of lignocellulosics more adapted to industrial biorefineries.

## MATERIALS AND METHODS

### Production of Plant Materials

The *proAtC4H::BdPMT1* construct used for poplar genetic transformation was the same as the one described in (Sibout et al., 2016), with the *BdPMT1* sequence inserted into the pCC0996 vector under the control of the *AtC4H* promoter (Weng et al., 2008). This construct was introduced using *A. tumefaciens* cocultivation into the hybrid poplar (*P. tremula x P. alba*) clone INRA 717-1B4 as well as in a 717-1B4 transgenic line named *AtF5H-OE*, according to the method described in Leplé et al., (1992). The *AtF5H-OE* line was previously transformed with an *AtF5H* gene under the control of the promoter of the *P. tremula x P. alba CesA4* gene (Potri.002G257900) (*proPtaCesA4::AtF5H).* Several transgenic lines from both genetic background were selected for further analyses. Two to five ramets of each line were acclimatized and grown in a S2 greenhouse for 3 months, from April until July. Height and stem diameter were measured before plant sampling for molecular and biochemical analyses.

Differentiating xylem samples were collected by a light scraping at the surface of the debarked stem. Samples were immediately frozen in liquid nitrogen and stored at −80°C until use. DNA was prepared using Nucleospin DNA Plant II kit (Macherey-Nagel, Hoerdt, France) and the integration of *BdPMT* and *F5H* genes was verified by PCR using the following primers pairs: PMT 5’-CCTCATCATGCAGGTGACAG-3’ and 5’-GAAGCAGTTGCCGTAGAACC-3’; F5H 5’-ATGGAGTCTTCTATATCACA-3’ and 5’-TTAAAGAGCACAGATGAGGC-3’. Likewise, RNA was extracted from differentiating xylem using a Nucleospin RNA Plant kit (Macherey-Nagel, Hoerdt, France). The expression level of the *BdPMT1* and *AtF5H* gene in each tree was evaluated by semi-quantitative RT-PCR performed in standard conditions using the same primers as above.

### Analyses of Cell Wall Phenolics

#### Preparation of CW Samples and Dioxane Lignins

All the analyses of cell wall phenolics were carried out from biological replicates (2, 3 or 4 per line) harvested from 3-month-old poplar trees. For each tree, the 20 cm-long basal part of the stem was collected, manually debarked, air-dried and ground to 0.5 mm. Extract-free samples were prepared by exhaustive water and ethanol extraction in an accelerated solvent extractor (ASE350, Dionex). The dried and extract-free samples are referred to as cell wall samples (CW).

The isolation of DL fractions was performed from 1 to 2 g of CW as previously described (Sibout et al., 2016). FTIR spectra of DL fractions were run on a Thermo Scientific Nicolet IS5 spectrophotometer and in KBr pellets.

#### Analytical Pyrolysis

Pyrolysis-gas chromatography-mass spectrometry (Py-GC/MS) was done using a CDS model 5250 pyroprobe autosampler interfaced to an Agilent 6890/5973 GC/MS. The CW samples (about 300 μg) were pyrolyzed in a quartz tube at 500°C for 15 s. The pyrolysis products were separated on a capillary column (5% phenyl methyl siloxane, 30 m, 250 μm i.d., and 0.25 μm film thickness) using helium as the carrier gas with a flow rate of 1 mL/min. The pyrolysis and GC/MS interfaces were kept at 290°C and the GC was programmed from 40°C (1 min) to 130°C at +6°C min^-1^, then from 130 to 250°C at +12°C min^-1^ and finally from 250°C to 300°C at +30°C min^-1^ (3 min at 300°C). The various phenolic pyrolysis compounds were identified by comparison to published spectra (Ralph and Hatfield, 1991). Py-GC/MS in the presence of tetramethylammonium hydroxyde (TMAH) was similarly performed but with addition of 3 μL of a 25% TMAH methanolic solution (Aldrich) onto the CW sample. The methylated pyrolysis products were identified by comparison of their mass spectra with those of the NIST MS library or with published TMAH-pyrograms (Kuroda et al., 2001; Kuroda et al., 2002).

#### Determination of Lignin Content

The determination of KL content was perfomed from about 300 mg of CW (weighted to the nearest 0.1 mg) and as previously described (Méchin et al., 2014). The quantitation of ABL was done from about 5 mg of CW (weighted to the nearest 0.01 mg) according to a recently published procedure (Sibout et al., 2016).

#### Determination of Ester-linked p-Hydroxybenzoic and p-Hydroxycinnamic Acids by Mild Alkaline Hydrolysis

About 5 to 10 mg of poplar CW or DL samples were put into 2-mL Eppendorf tube together with 1 mL of 1 M NaOH and 0.1 mL of *o*-coumaric internal standard (IS) methanolic solution. The IS amount was 0.05 mg for CW samples and 0.25 mg for DL ones. Mild alkaline hydrolysis was proceeded on a carousel overnight and at room temperature. After acidification (0.2 mL of 6 M HCl) and centrifugation (1500 g, 10 min), the supernatant was subjected to solid phase extraction as previously described (Ho-Yue-Kuang et al., 2016). The recovered methanolic samples were analyzed by HPLC combined with diode array detection (HPLC-DAD). For HPLC separation, 1 μL of sample was injected onto an RP18 column (4 × 50 mm, 2.7 μm particle size, Nucleoshell, Macherey-Nagel) with a flow rate of 0.25 mL min^−1^. The eluents were 0.1% formic acid in water (A) and 0.1% formic acid in acetonitrile (B), and the gradient was as follows: 0 min 5% B; 12 min, 20% B; 14 min, 80% B; 16 min, 5% B. The quantitative determination of alkali-released Bz, CA and FA was performed from the 250-400 nm DAD chromatograms and after calibration with authentic compounds

#### Analysis of Lignin Structure by Thioacidolysis

Thioacidolysis (4-hour long) followed by GC/MS of the trimethylsilylated (TMS) lignin-derived compounds was carried out from about 10 mg of CW samples using the simplified procedure previously published (Méchin et al., 2014), with some adaptations to the CW type concerning the IS amount and the reagent to sample ratio. In brief, 5 to 10 mg (weighed to the nearest 0.1 mg) were put together with 2 mL of freshly prepared thioacidolysis reagent and 0.1 mL of IS solution (heinecosane C21, 5 mg/mL in CH_2_Cl_2_) in a glass tube (Teflon-lined screwcap). The closed tubes were then heated at 100°C (oil bath) and for 4 h with occasional gentle shaking. After tube cooling, 2 mL of 0.2 M NaHCO_3_ were added to destroy the excess of BF_3_ etherate. Then, 0.025 mL of 6 M HCl were added to ensure that the pH was less than 3, before addition of 2 mL CH_2_Cl_2_ and tube mixing. A small amount (about 0.5 mL) of the lower organic phase was withdrawn with a glass Pasteur pipette, dried over anhydrous Na2SO4 and then directly subjected to trimethylsilylation. This silylation was performed with 10 μL of the solution together with 100 μL BSTFA (Sigma-Aldrich) and 10 μL of GC-grade pyridine (1 h at room temperature). The GC/MS analyses were carried out as previously described (Méchin et al., 2014). Some short thioacidolysis assays (1-hour long) were also carried out and were followed by desulfuration experiments according to a published method (Lapierre et al., 1995). In addition, thioacidolysis from exhaustively permethylated CW samples was run according to Sibout et al. (2016) and using the same thioacidolysis and GC/MS conditions.

### Investigation of Some CW Properties

#### Alkali Solubilization Assays

About 300 mg of poplar CW were subjected to mild alkaline hydrolysis in 10 mL of 1 M NaOH, into a 25 mL plastic tube agitated overnight on a carousel and at room temperature. The alkali-treated residue, referred to as the saponified residue (SR), was recovered by centrifugation (2000 g, 20 min), washed with 1 M HCl before centrifugation and then with water (3 times with centrifugation following each washing step). The final residue was freeze-dried, weighted to calculate its recovery yield and subjected to KL or ABL determination. The weight percentage of alkali-soluble lignin (% Alk-L) was calculated from the weight percentages of ABL in CW (%ABL_CW_) and in SR (%ABL_SR_) samples and from the SR recovery yield (%SR), as follows: % Alk-L = (100 x %ABL_CW_ – (%SR x %ABL_SR_)) / %ABL_CW_

#### Saccharification Assays

Saccharification experiments were performed from about 30 mg of SR samples (weighed to the nearest 0.1 mg) under the conditions previously described (Sibout et al., 2016). Saccharification efficency was calculated both from the weight loss and from the glucose yield.

## Acknowlegments

We thank Armelle Delile, Orlane Touzet (INRAE Biofora) and Anita Rinfray (GBFOR, INRAE, Forest Genetics and Biomass Facility, https://doi.org/10.15454/1.5572308287502317E12) for their technical contribution to produce and characterize the plant material. Likewise, the technical assistance of Frédéric Legée for the determination of Klason Lignin is gratefully acknowledged. The production of transgenic material has benefited from the LICA (Laboratoire d’Ingéniérie Cellulaire de l’Arbre) equipments.

## Supplemental Data

The following supplemental materials are available.

**Supplemental Figure S1.** Macroscopical and histochemical description of the plant material.

**Supplemental Figure S2.** IR spectra (KBr pellet) of DL lignin fractions isolated from *BdPMT1-OE/WT* and *BdPMT1-OE/AtF5H-OE* lines as compared to their controls.

**Supplemental Figure S3**. HPLC and GC/MS analyses of low-molecular weight phenolics released by alkaline hydrolysis of DL lignin fractions isolated from WT and *BdPMT1-OE/WT* lines.

**Supplemental Figure S4**. Correlation between the amount of ester-linked CA and the relative % of 4-vinylphenol (% VP) released by analytical pyrolysis of *BdPMT1* – OE poplar trees.

**Supplemental Figure S5**. Partial GC/MS chromatograms of the main dimers obtained after 1- or 4-hour-long thioacidolysis followed by Raney nickel desulfuration and from WT or *BdPMT1*-OE/WT lines.

**Supplemental Table S1.** Amount of *p*-coumaric acid (CA) ester-linked to grass CW and to the corresponding purified DL fractions.

**Supplemental Table S2.** Relative importance (% area) of the main dimers obtained after thioacidolysis and Raney nickel desulfuration of extract-free poplar stems.

## Notes

**Funding information:** the *AtF5H-OE* line was generated in the frame of the European ENERGYPOPLAR project (FP7-211917). The acquisition of the Py-GC/MS equipment was supported by the 3BCAR Carnot Institute.

